# Build a Better Bootstrap and the RAWR Shall Beat a Random Path to Your Door: Phylogenetic Support Estimation Revisited

**DOI:** 10.1101/2020.02.02.931063

**Authors:** Wei Wang, Kevin J. Liu

## Abstract

**Motivation:** The standard bootstrap method is used throughout science and engineering to perform general-purpose non-parametric resampling and re-estimation. Among the most widely cited and widely used such applications is the phylogenetic bootstrap method, which Felsenstein proposed in 1985 as a means to place statistical confidence intervals on an estimated phylogeny (or estimate “phylogenetic support”). A key simplifying assumption of the bootstrap method is that input data are independent and identically distributed (i.i.d.). However, the i.i.d. assumption is an over-simplification for biomolecular sequence analysis, as Felsenstein noted. Special-purpose fully parametric or semi-parametric methods for phylogenetic support estimation have since been introduced, some of which are intended to address this concern.

**Results:** In this study, we introduce a new sequence-aware non-parametric resampling technique, which we refer to as RAWR (“RAndom Walk Resampling”). RAWR consists of random walks that synthesize and extend the standard bootstrap method and the “mirrored inputs” idea of Landan and Graur. We apply RAWR to the task of phylogenetic support estimation. RAWR’s performance is compared to the state of the art using synthetic and empirical data that span a range of dataset sizes and evolutionary divergence. We show that RAWR support estimates offer comparable or typically superior type I and type II error compared to phylogenetic bootstrap support as well as GUIDANCE2, a state-of-the-art purpose-built fully parametric method. Additional simulation study experiments help to clarify practical considerations regarding RAWR support estimation. We conclude with thoughts on future research directions and the untapped potential for sequence-aware non-parametric resampling and re-estimation.

**Availability:** Data and software are publicly available under open-source software and open data licenses at: https://gitlab.msu.edu/liulab/RAWR-study-datasets-and-scripts.

## 1 Introduction

In 1985, Felsenstein proposed the application of standard bootstrap resampling (Efron, 1979) to place confidence intervals on an estimated phylogeny (Felsenstein, 1985). Given an input multiple sequence alignment (MSA), the approach first generates bootstrap replicates by sampling input MSA columns uniformly at random with replacement. Then, phylogenetic re-estimation is performed on each bootstrap replicate. Finally, bootstrap support for each edge of an annotation phylogeny (i.e., the phylogeny estimated on the original input MSA) is calculated as the fraction of re-estimated phylogenies that also display that edge.

Bootstrap support estimation has become a de facto standard for assessing reproducibility in modern phylogenetics and phylogenomics, and Felsenstein’s seminal 1985 paper has become the 41st most cited in all of science, according to the 2014 survey of Van Noorden *et al.* (2014). Alternatives include other non-parametric resampling methods such as the jackknife (Tukey, 1958) and parametric resampling (Warnow, 2017; Felsenstein, 2004). Examples of the latter include MSA-specific confidence measures such as GUIDANCE1 (Landan and Graur, 2008; Penn *et al.*, 2010), GUIDANCE2 (Sela *et al.*, 2015), PSAR (Kim and Ma, 2011), T-COFFEE (Notredame *et al.*, 2000), wpSBOOT (Chang *et al.*, 2019), and Divvier (Ali *et al.*, 2019) and parametric and/or special-purpose MSA resampling or filtering techniques applied to the problem of phylogenetic support estimation, including TCS (Chang *et al.*, 2014), the unistrap (Chatzou *et al.*, 2018), Gblocks (Talavera and Castresana, 2007), Trimal (Capella-Gutiérrez *et al.*, 2009), and the method of Rajan (2013). Both classes of alternative methods are less popular than the bootstrap. Furthermore, parametric resampling/filtering methods require the assumption that data are generated from a particular parametric model, and special-purpose resampling/filtering methods do not readily generalize beyond a specific application. Our study focuses on non-parametric techniques for phylogenetic support estimation for these reasons.

But the bootstrap also makes a key simplifying assumption. Felsenstein concluded his landmark paper with a forward-thinking cautionary note:

> “A more serious difficulty is the lack of independence of the evolutionary processes in different characters…. For the purposes of this paper, we will ignore these correlations and assume that they cause no problems; in practice, they pose the most serious challenge to the use of bootstrap methods.”

Crucially, a variety of biological factors violate the i.i.d. assumption, including evolutionary processes such as genetic recombination and sequence insertions/deletions.

We and several co-authors created the SERES (or “SEquential RESampling”) method to relax this simplifying assumption (Wang *et al.*, 2018). SERES generalizes the bootstrap and the “mirrored inputs” idea of Landan and Graur (2007) into a random walk on either unaligned or aligned biomolecular sequence inputs. A critical property of the random walk procedure is “neighbor preservation”: neighboring bases that appear in a resampled sequence are also guaranteed to be neighbors in the corresponding untransformed sequence in the original input. SERES resampling of unaligned sequences requires synchronization points in the form of anchor regions, similar to the use of barriers in asynchronous computing. To date, the utility of SERES resampling and re-estimation has been demonstrated on two applications – placing confidence intervals on estimated multiple sequence alignments (Wang *et al.*, 2018) and temporal model inference and learning (Wang *et al.*, 2019; Wuyun *et al.*, 2019).

## 2 Approach

In this study, we return to where we started: Felsenstein’s landmark 1985 contribution – the phylogenetic bootstrap support method. We now briefly summarize the contributions of our study. Our approach to the problem of phylogenetic support estimation makes use of a simpler resampling strategy which we refer to as RAWR (or “RAndom Walk Resampling”). The relative simplicity of RAWR compared to SERES, our previous method for semi-parametric resampling, is the first contribution of our study. RAWR does not require a parametric method for anchor estimation, where as SERES does, adding complexity in the form of method parameters such as anchor width, count, and sequence conservation criterion used for selecting anchors. Second, our approach focuses on sequence-aware non-parametric resampling and re-estimation using unaligned sequence data, which differs from existing studies and widely-used state-of-the-art methods. Third, RAWR phylogenetic support estimates offer similar or typically better accuracy compared to the bootstrap method – the de facto standard for phylogenetic and phylogenomic confidence interval estimation – as well as a leading special-purpose parametric resampling method.

## 3 Methods

The computational problem addressed in our study is defined as follows. The problem inputs consist of an MSA A that was estimated using an MSA method *f* and a phylogenetic tree *T* that was estimated using a phylogenetic inference method *g* applied to *A*. The problem output consists of confidence interval estimates (or support estimates) *ϵ*(*e*) ∈ [0,1] for every non-leaf edge *e* ∈ *T*.

### RAWR

The RAWR method for phylogenetic support estimation conducts a random walk on the input MSA *A*. Each random walk produces a resampled replicate set of unaligned sequences, and the resampling procedure is repeated to obtain a set of resampled replicates. MSA and tree re-estimation is then performed on the resampled replicates. Phylogenetic support for a branch in the original estimated tree is calculated to be the fraction of re-estimated trees that also display the branch. Pseudocode for RAWR phylogenetic support estimation is provided in Figure 1. Our performance study utilized reversal probability *γ* ∈ {1 × 10^−3^, 1 × 10^−2^, 1 ×10^−1^, 2×10^−1^, 3×10^−1^}. The default choice for *γ* in our study was 1 ×10^−1^ unless otherwise noted. RAWR was used to generate 100 resampled replicates for each input dataset.

**Fig. 1.**
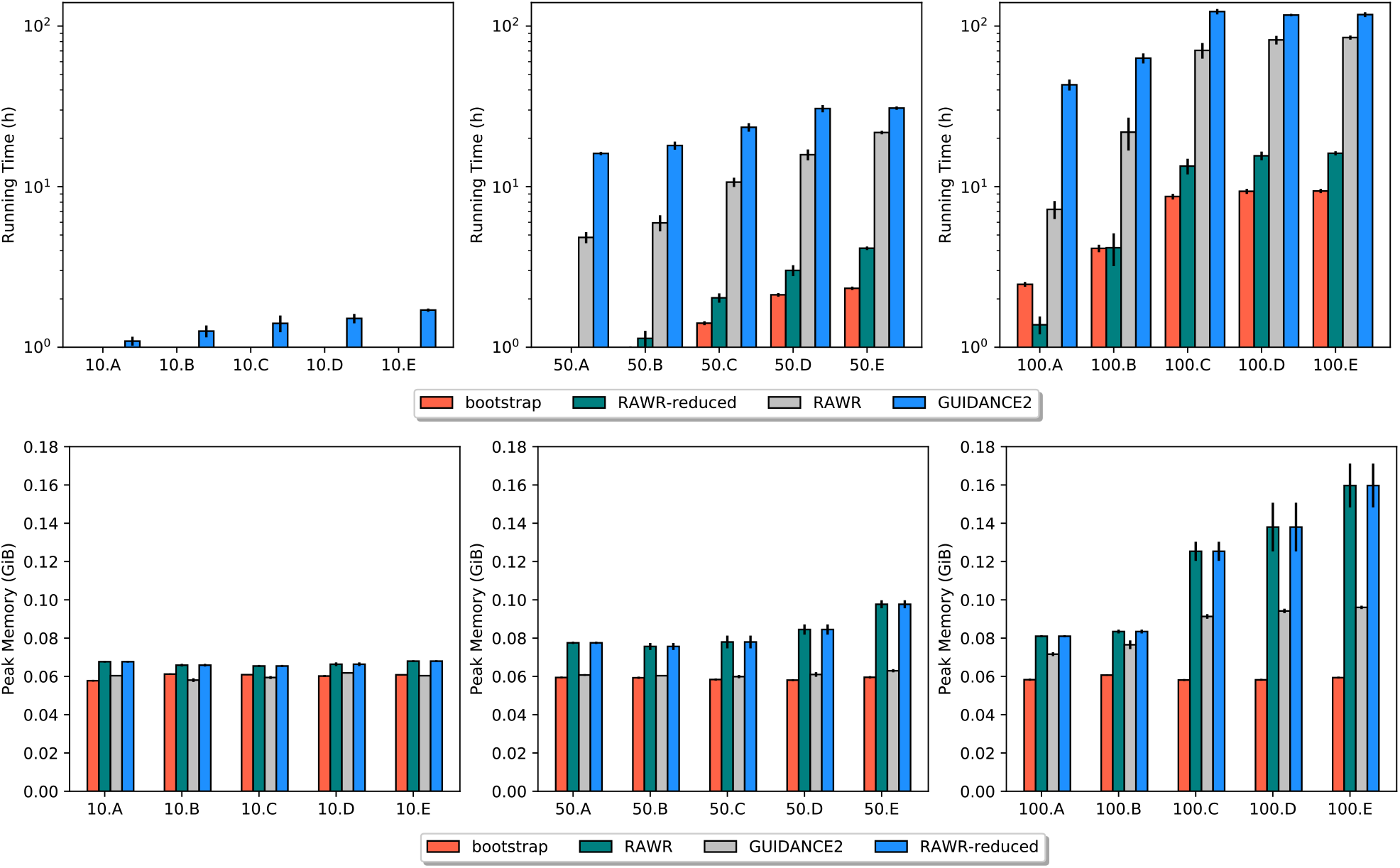
Simulation study: runtime and memory usage comparison of phylogenetic support estimation methods. Results are shown for the 10-taxon, 50-taxon, and 100-taxon model conditions in the left, middle, and right columns, respectively. The 10-taxon model conditions are arranged on the x-axis from left to right in order of generally increasing evolutionary divergence; the 50-taxon and 100-taxon model conditions are similarly arranged. In each panel, average serial runtime or peak memory usage across all replicate datasets in the model condition is shown along with standard error bars (*n* = 20). In the top row of panels, the y-axis shows serial runtime in hours and is in log-scale. In the bottom row of panels, the y-axis shows peak memory usage in GiB and is in absolute scale. Also note that the figure legends differ between the top and bottom row of panels.

### Methods for MSA and phylogenetic estimation/re-estimation

We focused on two-phase methods for phylogenetic inference on unaligned DNA sequences. This class of methods is by far the most prevalent in systematic studies. The first phase of a two-phase method estimates a multiple sequence alignment on an input set of unaligned sequences, and the second phase uses the previous phase’s estimated MSA to estimate a phylogenetic tree. Our performance study included MSA methods that varied in terms of alignment accuracy and computational efficiency. MAFFT is a popular suite of MSA algorithms which has been shown to be among the most accurate methods for both MSA estimation (Liu *et al.*, 2009, 2012) and, when used in combination with leading MLE phylogenetic inference methods, phylogenetic inference from unaligned sequences (Liu *et al.*, 2009, 2012) We ran version 7.222 of MAFFT software with default settings, based on the following command:

~~~
mafft <sequence file>
  > <estimated alignment file>
~~~

To explore the impact of alignment quality on downstream phylogenetic inference and support estimation, we also included ClustalW since it was among the first popular MSA methods and has become a mainstay throughout computational biology and bioinformatics. Our ClustalW analyses were run using the version 2.1 software with default settings, as specified by the following command:

~~~
clustalw2 -INFILE=<resampled sequence file>
  -ALIGN -TYPE=dna
  -outfile=<estimated alignment file>
  -output=FASTA
~~~

Summary statistics for the estimated MSAs are shown in Tables 1 and 2. Maximum likelihood phylogenies under the GTR+Γ model (Rodriguez *et al.*, 1990; Yang, 1993; Wakeley, 1993) were inferred on estimated MSAs using RAxML (Stamatakis, 2014). We used version 8.2.11 of the RAxML software. RAxML analyses (other than bootstrap analyses) were performed using the following command:

**Table 1.** Simulation study: model condition parameters and summary statistics. Model condition parameters consisted of the number of taxa, tree height, and insertion/deletion probability. Each 10-taxon model condition is named 10.A through 10.E in generally increasing order of evolutionary divergence; the 50-taxon and 100-taxon model conditions are named similarly. The following average summary statistics are reported for true MSAs and MAFFT-estimated MSAs on each model condition (*n* = 20): “ANHD” is the average normalized Hamming distance of a pair of aligned sequences in an MSA, “Gappiness” is the proportion of an MSA matrix that consists of indels, and “length” is the number of MSA columns, and “SP-FN” and “SP-FP” are the proportions of nucleotide-nucleotide homologies that appear in the true alignment but not in the estimated alignment or vice versa, respectively. The average normalized Robinson-Foulds distance (“nRF”) between the model tree and the RAxML(MAFFT)-inferred tree is also reported for each model condition (*n* = 20).

| Model condition | Number of taxa | Model Tree height | Insertion/deletion probability | True alignment |  |  | MAFFT alignment |  |  | RAxML (MAFFT) nRF |
| --- | --- | --- | --- | --- | --- | --- | --- | --- | --- | --- |
|  |  |  |  | ANHD | Gappiness | length | length | SP-FN | SP-FP |  |
| 10.A | 10 | 0.47 | 0.13 | 0.380 | 0.591 | 2466 | 1543 | 0.566 | 0.629 | 0.186 |
| 10.B | 10 | 0.7 | 0.1 | 0.479 | 0.618 | 2691 | 1602 | 0.687 | 0.750 | 0.243 |
| 10.C | 10 | 1.2 | 0.06 | 0.591 | 0.645 | 2832 | 1588 | 0.811 | 0.850 | 0.443 |
| 10.D | 10 | 2 | 0.031 | 0.642 | 0.591 | 2490 | 1583 | 0.815 | 0.841 | 0.464 |
| 10.E | 10 | 4.4 | 0.013 | 0.696 | 0.578 | 2390 | 1623 | 0.904 | 0.913 | 0.664 |
| 50.A | 50 | 0.45 | 0.06 | 0.415 | 0.667 | 3070 | 2053 | 0.340 | 0.336 | 0.084 |
| 50.B | 50 | 0.73 | 0.03 | 0.513 | 0.603 | 2525 | 1834 | 0.451 | 0.431 | 0.146 |
| 50.C | 50 | 1.2 | 0.02 | 0.598 | 0.620 | 2646 | 1950 | 0.731 | 0.704 | 0.322 |
| 50.D | 50 | 2 | 0.012 | 0.667 | 0.629 | 2720 | 2171 | 0.902 | 0.881 | 0.517 |
| 50.E | 50 | 4.3 | 0.005 | 0.715 | 0.591 | 2474 | 2385 | 0.974 | 0.965 | 0.755 |
| 100.A | 100 | 4 | $1 \times 10^{-5}$ | 0.454 | 0.331 | 1682 | 1533 | 0.054 | 0.046 | 0.075 |
| 100.B | 100 | 7 | $1 \times 10^{-5}$ | 0.540 | 0.439 | 2263 | 1861 | 0.209 | 0.176 | 0.119 |
| 100.C | 100 | 15 | $5 \times 10^{-5}$ | 0.646 | 0.571 | 2317 | 2418 | 0.680 | 0.603 | 0.470 |
| 100.D | 100 | 25 | $2 \times 10^{-5}$ | 0.683 | 0.634 | 1837 | 2799 | 0.899 | 0.853 | 0.607 |
| 100.E | 100 | 20 | $4 \times 10^{-5}$ | 0.672 | 0.614 | 2487 | 2701 | 0.848 | 0.796 | 0.661 |

**Table 2.**
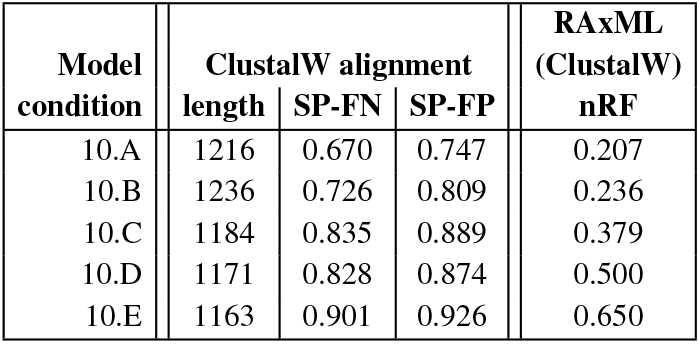
Simulation study: summary statistics for ClustalW alignments and RAxML(ClustalW) trees on 10-taxon model conditions. Table layout and description are otherwise identical to Table 1.

~~~
raxmlHPC -s <estimated alignment file>
  -n <name> -m GTRGAMMA -p <random number>
  -# 10
~~~

#### Algorithm 1 RAWR phylogenetic support estimation

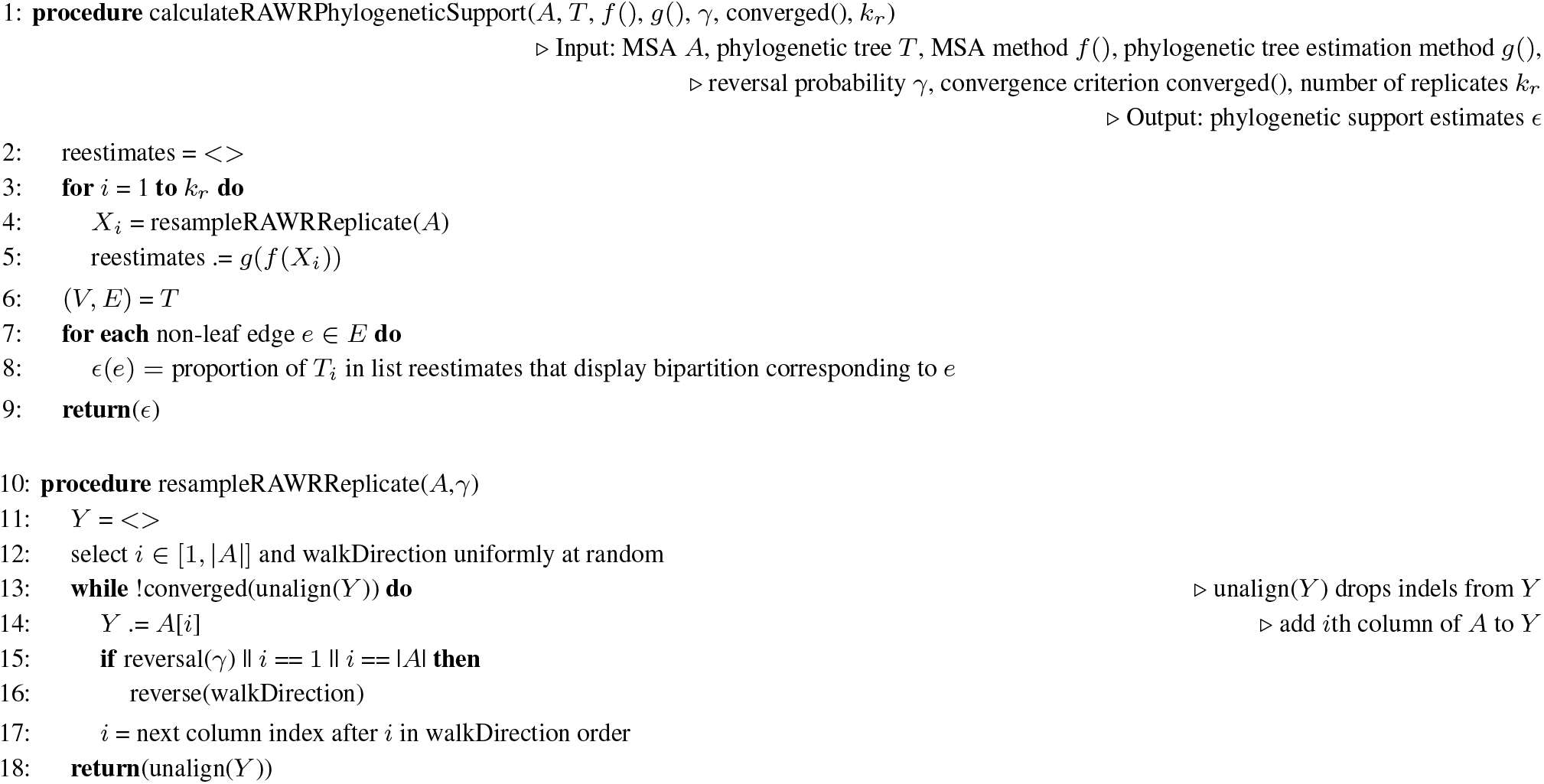

For brevity, RAxML(*x*) denotes a two-phase analysis consisting of running method *x* to first estimate an MSA on an input set of unaligned sequences, followed by running RAxML on the *x*-estimated MSA as input.

### Phylogenetic bootstrap method

Bootstrap analyses were run using the bootstrap implementation in version 8.2.11 of the RAxML software. The analyses were run using the following command:

~~~
raxmlHPC -s <estimated alignment file>
  -n <name> -m GTRGAMMA
  -p <random number> -b <random number>
  -# <number of sample tree>
~~~

Each bootstrap support analysis utilized 100 bootstrap replicates; downstream re-estimation and support calculation steps were otherwise identical to RAWR.

### GUIDANCE2

GUIDANCE2 augments Landan and Graur (2007)’s “mirrored inputs” idea with progressive MSA-specific parametric resampling techniques. GUIDANCE2 was run with default settings. For each dataset, GUIDANCE2 was used to resample 100 replicates and reestimation was performed on each resampled replicate using an identical procedure as in the RAWR and bootstrap analyses (see above); the support calculation made use of the RAxML software and the following command:

~~~
raxmlHPC -f b -m GTRGAMMA
  -t <inferred tree file>
  -z <sampled trees file>
  -n <name>
~~~

Although Landan and Graur (2007) originally focused on MSA estimation and GUIDANCE2 was originally developed for MSA confidence interval placement, it is natural to consider the impact of MSA quality on downstream inference tasks. As we demonstrate in our performance study, a new application of these parametric and semi-parametric techniques beyond their originally intended use can bring value.

### Simulation study datasets

Our simulation study utilized model conditions and synthetic benchmarks from two previous studies (Wang *et al.*, 2018; Liu *et al.*, 2012). The model conditions exhibit varying dataset sizes and evolutionary divergence that are meant to capture a range of computational difficulty. We briefly recap the experimental procedures used for simulation. First, model trees were sampled as follows. For the 10-taxon and 50-taxon simulations, INDELible version 1.03 (Fletcher and Yang, 2009) was used to sample non-ultrametric trees under a random birth process with branch lengths drawn uniformly at random from the open unit interval. For the 100-taxon simulations, random birth-death model trees were sampled using r8s version 1.7 (Sanderson, 2003) and the following script:

~~~
begin r8s;
simulate diversemodel=bdback seed=<random seed>
  nreps=20 ntaxa=<10 or 50> T=0;
describe tree=0 plot=chrono_description;
end;
~~~

The 100-taxon model trees were then deviated from ultrametricity using Nakhleh *et al.* (2002)’s procedure with deviation factor *c* = 2.0; the procedure was performed using a custom script (available at http://www.cs.utexas.edu/users/tandy/science-paper.html and https://github.com/tandyw/datasets/). Model trees were then rescaled to obtain total height specified by model condition parameter *h*. Then, nucleotide sequence evolution on model trees was simulated under a finite-sites models of nucleotide substitutions and sequence insertions/deletions with root sequence length of 1 kb. The former consisted of the general time-reversible (GTR) model (Rodriguez *et al.*, 1990) with base frequency and substitution rate parameter settings based upon empirical NemAToL estimates from the study of Liu *et al.* (2012). INDELible (Fletcher and Yang, 2009) was used to simulate 10- and 50-taxon datasets under the GTR model and the indel model of (Fletcher and Yang, 2009). The 100- taxon simulations were performed using ROSE under the GTR model and an indel model with a “medium” gap length distribution, as described by the earlier study of (Liu *et al.*, 2012). For each model condition, the simulation procedure was repeated to obtain 20 experimental replicates. Model condition parameters and summary statistics for simulated datasets are listed in Table 1.

### Empirical study datasets

The empirical benchmarking data used in our study was obtained from the Comparative RNA Website (CRW) database (accessible at www.rna.icmb.utexas.edu) (Cannone *et al.*, 2002). Almost all of the CRW rRNA datasets are provided with comprehensively curated multiple sequence alignments that were produced using intensive hybrid (i.e., automated and human) analysis of biomolecular sequence, structural, and other information; the curated sequence alignments represent a “gold standard” reference for benchmarking studies involving sequence alignment tasks (Liu *et al.*, 2009, 2012). The reference alignments were used to obtain reference trees, which consisted of MLE trees estimated on reference alignment. RAxML was used to perform these analyses using the same command as in the simulation study. As in earlier studies (Liu *et al.*, 2009, 2012; Mirarab *et al.*, 2015), our choice of reference tree is a practical one in the absence of ground truth.

Our simulation study model conditions best reflect non-coding nucleotide sequence evolution, and our empirical study focuses on intronic rRNA datasets for experimental consistency. Similar to the simulation study, we selected datasets with a range of evolutionary divergence and dataset sizes up to 250 sequences. Sequences with greater than 99% missing data were omitted from analysis. Table 3 provides summary statistics and other information on the empirical datasets.

**Table 3.** Empirical study: summary statistics. The empirical datasets used in our performance study were obtained from the Comparative RNA Website (CRW) database (Cannone *et al.*, 2002). (See Methods section for more details.) The curated alignment and a maximum likelihood tree estimated on the curated alignment were used as the reference alignment and tree for benchmarking purposes. We report the following summary statistics for each reference alignment (*n* = 1): “ANHD” or average normalized Hamming distance, “Gappiness”, and “length”; we also report the MAFFT-estimated MSA “length” and “SP-FN” and “SP-FP” errors with respect to the reference alignment, as well as the “nRF” or normalized Robinson-Foulds distance between the RAxML(MAFFT) tree and the reference tree. Summary statistic calculations and descriptions are otherwise identical to Table 1.

| Dataset | Number of taxa | Reference alignment |  |  | MAFFT alignment |  |  | RAxML (MAFFT) nRF |
| --- | --- | --- | --- | --- | --- | --- | --- | --- |
|  |  | ANHD | Gappiness | length | length | SP-FN | SP-FP |  |
| IGIA | 110 | 0.606 | 0.915 | 10368 | 6065 | 0.732 | 0.780 | 0.645 |
| IGIB | 202 | 0.579 | 0.910 | 10633 | 7070 | 0.825 | 0.863 | 0.678 |
| IGIC2 | 32 | 0.533 | 0.700 | 4243 | 3530 | 0.691 | 0.716 | 0.517 |
| IGID | 21 | 0.719 | 0.782 | 5061 | 3063 | 0.874 | 0.905 | 0.778 |
| IGIE | 249 | 0.451 | 0.838 | 2751 | 2847 | 0.406 | 0.389 | 0.585 |
| IGIIA | 174 | 0.668 | 0.814 | 6406 | 6945 | 0.817 | 0.800 | 0.450 |

### Performance criteria

Phylogenetic support estimation methods were evaluated based on type I and type II error of estimated phylogenetic support for a phylogenetic tree estimate with respect to a reference tree (i.e., the model tree for each simulated dataset and the reference tree for each empirical dataset). We used precision-recall (PR) curves and area under PR curves (PR-AUC) to evaluate both types of error and tradeoffs between them. We note that the phylogenetic support estimation problem (and, more generally, classical estimation of confidence intervals in statistics) requires an original estimate for annotation; a two-phase method was used for this purpose (see “Methods for MSA and phylogenetic estimation/re-estimation” text above). A phylogenetic support estimation method was then run to annotate each branch of the estimated tree with a phylogenetic support value and thereby place confidence intervals on the estimated tree topology. For this reason, the confusion matrix for the ROC and PR curves was formed from the following four classes: true positives (TP) consist of bipartitions of the estimated tree that have support value greater than or equal to a given threshold and appear in the reference tree, false positives (FP) consist of bipartitions of the estimated tree that have support value greater than or equal to a given threshold but do not appear in the reference tree, false negatives (FN) consist of bipartitions of the estimated tree that have support less than a given threshold but appear in the reference tree, and true negatives (TN) consist of bipartitions of the estimated tree that have support less than a given threshold and do not appear in the reference tree.

The PR curve plots true positive rate 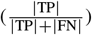 vs. precision 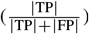, and varying the threshold for confusion matrix calculations yields different points along the curve. We used custom scripts and the scikit-learn Python library (Pedregosa *et al.*, 2011) to calculate the curves and AUC quantities.

We also compared phylogenetic support estimation methods based on serial computational runtime and peak main memory usage. All experiments were conducted on computing facilities in the Michigan State University High Performance Computing Center. We used compute nodes in the intel16-k80 cluster, each of which had a 2.4 GHz 14-core Intel Xeon E5-2680v4 processor.

## 4 Results

### 4.1 Simulation Study

#### Performance comparison of RAWR versus bootstrap and GUIDANCE2

We compared the performance of RAWR versus bootstrap and GUIDANCE2 on the simulation study datasets, where MAFFT and/or RAxML(MAFFT) were used to estimate/re-estimate MSAs and phylogenetic trees. The type I and type II error of the different methods were evaluated based on PR-AUC, as shown in Table 4.

**Table 4.**
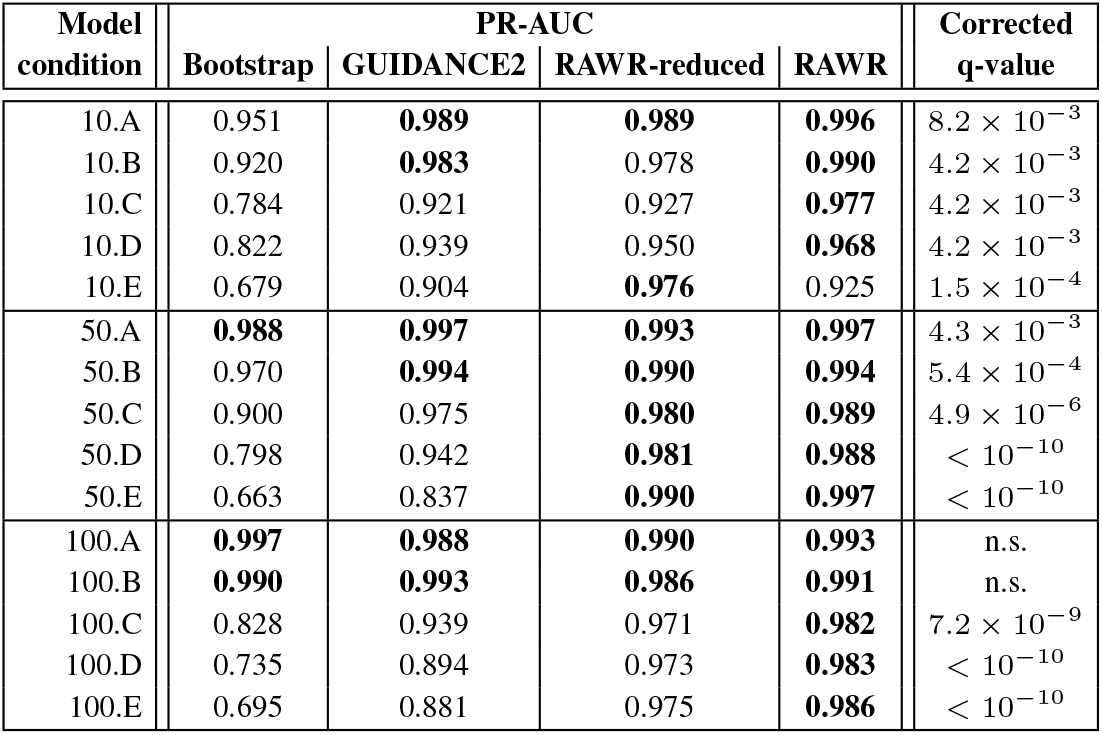
Simulation study: PR-AUC comparison of phylogenetic support estimation methods. MAFFT and RAxML(MAFFT) were used to perform MSA and tree estimation/re-estimation, respectively. We report each method’s aggregate PR-AUC across all replicate datasets for a model condition (*n* = 20). For each model condition, the top PR-AUC values within an absolute difference of 0.01 are shown in bold. Statistical significance of PR-AUC differences between RAWR and bootstrap were evaluated using a one-tailed pairwise t-test and a multiple test correction was performed using the method of Benjamini and Hochberg (1995). Corrected q-values are reported (*n* = 20).

| Model condition | PR-AUC |  |  |  | Corrected q-value |
| --- | --- | --- | --- | --- | --- |
|  | Bootstrap | GUIDANCE2 | RAWR-reduced | RAWR |  |
| 10.A | 0.951 | **0.989** | **0.989** | **0.996** | $8.2 \times 10^{-3}$ |
| 10.B | 0.920 | **0.983** | 0.978 | **0.990** | $4.2 \times 10^{-3}$ |
| 10.C | 0.784 | 0.921 | 0.927 | **0.977** | $4.2 \times 10^{-3}$ |
| 10.D | 0.822 | 0.939 | 0.950 | **0.968** | $4.2 \times 10^{-3}$ |
| 10.E | 0.679 | 0.904 | **0.976** | 0.925 | $1.5 \times 10^{-4}$ |
| 50.A | **0.988** | **0.997** | **0.993** | **0.997** | $4.3 \times 10^{-3}$ |
| 50.B | 0.970 | **0.994** | **0.990** | **0.994** | $5.4 \times 10^{-4}$ |
| 50.C | 0.900 | 0.975 | **0.980** | **0.989** | $4.9 \times 10^{-6}$ |
| 50.D | 0.798 | 0.942 | **0.981** | **0.988** | $< 10^{-10}$ |
| 50.E | 0.663 | 0.837 | **0.990** | **0.997** | $< 10^{-10}$ |
| 100.A | **0.997** | **0.988** | **0.990** | **0.993** | n.s. |
| 100.B | **0.990** | **0.993** | **0.986** | **0.991** | n.s. |
| 100.C | 0.828 | 0.939 | 0.971 | **0.982** | $7.2 \times 10^{-9}$ |
| 100.D | 0.735 | 0.894 | 0.973 | **0.983** | $< 10^{-10}$ |
| 100.E | 0.695 | 0.881 | 0.975 | **0.986** | $< 10^{-10}$ |

**Table 5.** Simulation study: RAWR support estimation using alternative estimation/re-estimation methods. We compared RAWR support estimation using two different estimation/re-estimation methods: either MAFFT and RAxML(MAFFT) or ClustalW and RAxML(ClustalW). For each of the two methods, aggregate PR-AUC is shown across all replicate datasets of each model condition (*n* = 20).

| Model condition | RAWR using |  |
| --- | --- | --- |
|  | MAFFT and RAXML(MAFFT) | ClustalW and RAXML(ClustalW) |
| 10.A | 0.996 | 0.997 |
| 10.B | 0.990 | 0.991 |
| 10.C | 0.977 | 0.988 |
| 10.D | 0.968 | 0.992 |
| 10.E | 0.925 | 0.964 |

**Table 6.** Simulation study: RAWR support estimation using different choices for reversal probability *γ*. Aggregate PR-AUC is reported across all replicate datasets of each 10-taxon model condition (*n* = 20).

| Model condition | Reversal probability $\gamma$ | | | | |
| --- | --- | --- | --- | --- | --- |
| | $1 \times 10^{-3}$ | $1 \times 10^{-2}$ | $1 \times 10^{-1}$ | $2 \times 10^{-1}$ | $3 \times 10^{-1}$ |
| 10.A | **0.997** | **0.998** | **0.994** | 0.986 | 0.980 |
| 10.B | **0.994** | **0.990** | **0.987** | **0.985** | 0.977 |
| 10.C | 0.942 | 0.941 | **0.968** | **0.957** | 0.901 |
| 10.D | **0.977** | **0.982** | 0.944 | 0.935 | 0.934 |
| 10.E | **0.969** | **0.978** | 0.929 | 0.923 | 0.922 |

RAWR consistently returned comparable or better PR-AUC compared to bootstrap and GUIDANCE2; RAWR improvements over bootstrap were statistically significant using pairwise t-tests with Benjamini Hochberg correction (Benjamini and Hochberg, 1995) (*n* = 20 and *α* = 0.05) on all model conditions with just two exceptions – the two least divergent 100- taxon model conditions, where all methods returned comparable PR-AUC (within 0.01 of the best method).

RAWR’s PR-AUC advantage over bootstrap and GUIDANCE2 tended to grow as model conditions grew larger and/or more divergent. This suggests that RAWR support estimates offers better type I and type II error on more challenging datasets. The maximum absolute PR-AUC improvements of RAWR over bootstrap and GUIDANCE2 were 0.334 and 0.160, respectively, and the averages across all model conditions were 0.136 and 0.039, respectively. A difference between the methods is that GUIDANCE2 and RAWR incorporate re-estimation of both MSAs and phylogenetic trees, but bootstrap only incorporates re-estimation of trees. This is because it doesn’t make sense to re-estimate an MSA on a bootstrap replicate, since the lack of neighbor preservation means a possible loss of sequence homology.

We note that GUIDANCE2 incorporates standard bootstrap resampling as a first step, and subsequent steps focus on guide tree re-estimation and other re-estimation tasks as part of progressive MSA re-estimation. For this reason, GUIDANCE2 can be seen as an adaptation of the standard bootstrap to MSA and tree re-estimation.

Finally, we note that GUIDANCE2 is purpose-built for MSA reestimation, whereas bootstrap and RAWR are general purpose nonparametric resampling methods (inasmuch as both resample an MSA without utilizing an explicit parametric model). Despite this, RAWR was able to match or exceed GUIDANCE2’s PR-AUC performance.

A performance comparison based on type I and type II error alone is incomplete, however. We also compared serial runtime and peak memory usage for all methods (Figure 1).

Compared to the bootstrap method, the non-bootstrap methods require an additional alignment re-estimation step on each resampled replicate. This key difference resulted in multiple factors of greater runtime for the former as compared to the latter, where all methods utilized the same amount of resampling replication (i.e., 100 replicates). GUIDANCE2 was the slowest method overall – even moreso than RAWR – due to the complexity of its special-purpose MSA re-estimation approach. Overall, absolute runtimes were relatively short on 10-taxon model conditions – on the order of minutes for bootstrap and RAWR – but grew quickly as the number of taxa increased. On the most divergent 100-taxon model condition, analyses took between half a day and multiple days to finish. We attribute observed runtime trends to the computational complexity of MSA estimation and phylogenetic MLE problems (Wang and Jiang, 1994; Roch, 2006).

A comparison of peak memory usage resulted in qualitatively similar method rankings, although the absolute differences were not as large as in the case of the runtime comparisons. Peak memory usage was modest in our study – amounting to just a few hundred MiB, an amount well within the scope of modern PC specifications. The bootstrap method typically utilized the least main memory, followed by GUIDANCE2 and then RAWR. As in other related studies (Liu *et al.*, 2012; Mirarab *et al.*, 2015), we anticipate that, due to the computational difficulty of the applications in our study, memory limitations will quickly become a major bottleneck as dataset sizes increase.

Overall, the simulation study experiments suggest that RAWR offers a performance advantage in terms of type I and type II error vs. the state of the art; this improvement come at the cost of increased time relative to a standard bootstrap analysis (but not a purpose-built parametric resampling method).

#### RAWR support estimation using reduced resampling replication

Our performance study also included a conservative comparison where we reduced the number of resampled replicates in our RAWR methods’ analyses by an order of magnitude (i.e., 10 resampled replicates as opposed to the 100 resampled replicated used by the other phylogenetic support estimation methods). We refer to the resulting method as “RAWR-reduced”. Despite the reduction of resampled data, RAWR- reduced returned comparable or better PR-AUC compared to bootstrap and GUIDANCE2, with greater PR-AUC improvements occurring on larger and/or more divergent model conditions. The PR-AUC returned by RAWR-reduced was comparable to RAWR, amounting to an average absolute difference of 0.007 across all model conditions. On average for each model condition, the RAWR-reduced analyses had slightly larger serial runtime compared to the bootstrap method, but were smaller than RAWR and GUIDANCE2 by multiple factors; both RAWR and RAWR- reduced had essentially identical peak memory usage. Thus, where computational efficiency is a concern, the use of reduced resampling replication in RAWR analyses allows a tradeoff between type I/II error and computational efficiency, without too great of a penalty for the former.

#### RAWR support estimation using alternative MSA/tree estimation/re-estimation methods

As in other performance studies of MSA and phylogenetic tree estimation from unaligned sequence inputs (Liu *et al.*, 2009, 2012), we found that MAFFT generally produced more accurate alignments than ClustalW on the 10-taxon model conditions, although this accuracy improvement did not translate directly to more accurate downstream phylogenetic inference (Tables 1 and 2).

Despite this, RAWR returned comparable PR-AUC regardless of which of the two MSA methods were used on the 10.A and 10.B model conditions. On the more divergent 10.C through 10.E model conditions, RAWR returned respective PR-AUC improvements of 0.011, 0.024, and 0.039 when using ClustalW and RAxML(ClustalW) for estimation/re-estimation, rather than MAFFT and RAxML(MAFFT). Our finding suggests that RAWR support estimation is robust to annotation MSA/tree quality, and hints at an even stronger result: neighbor-preserving random walks may yield better support estimates where computational problems are more difficult and estimation uncertainty is therefore greater.

#### RAWR support estimation using alternative choices for reversal probability *γ*

On each 10-taxon model condition except for the 10.C model condition, RAWR returned similar PR-AUC as the reversal probability *γ* was increased from 0.001 up until a critical threshold; PR-AUC then dropped as *γ* increased past the threshold. The exact threshold varied somewhat across model conditions. More generally, we observed a range of RAWR *γ* settings that returned the highest PR-AUC, where the range typically spanned around one to two orders of magnitude.

### 4.2 Empirical Study

We also evaluated RAWR versus bootstrap and GUIDANCE2 based on their PR-AUC on the empirical datasets (Table 7). The PR-AUC comparison outcomes were broadly similar to those seen in the simulation study. RAWR outperformed the bootstrap method on all empirical benchmarks, with an average absolute improvement of 0.105. RAWR also outperformed GUIDANCE2 on all empirical benchmarks except for IGIB. The average absolute difference of the two methods’ PR-AUC values was 0.055. For all three methods, the worst PR-AUC values in our entire study were observed on the IGIB dataset. One primary factor for this outcome is the extremely high gappiness of the reference alignments for the IGIB and IGIA datasets (i.e., the fraction of the reference alignment that consists of indels), as compared to every other dataset in our study. RAWR resampling of datasets with extremely high gappiness may require additional safeguards to mitigate the “de-synchronization” issues noted above.

**Table 7.** Empirical study: PR-AUC comparison of phylogenetic support estimation methods. MAFFT and RAxML(MAFFT) were used to perform MSA and tree estimation/re-estimation, respectively. For each empirical dataset, the top PR-AUC value is shown in bold.

| Dataset | PR-AUC |  |  |
| --- | --- | --- | --- |
|  | Bootstrap | GUIDANCE2 | RAWR |
| IGIA | 0.725 | 0.705 | **0.804** |
| IGIB | 0.629 | **0.737** | 0.695 |
| IGIC2 | 0.778 | 0.874 | **0.957** |
| IGID | 0.670 | 0.740 | **0.884** |
| IGIE | 0.772 | 0.777 | **0.808** |
| IGIIA | 0.830 | 0.870 | **0.884** |

Three differences between the empirical study and simulation study are worth noting. Reference trees in the empirical study are not the same as ground truth in the simulation study. The former consisted of MLE- inferred trees on highly accurate curated alignments, which in turn are expected to be highly accurate (but not perfectly accurate). Also, by necessity, the number of benchmarking datasets differed between the simulation and empirical study. This is due to the large amount of effort required to curate reference alignments for empirical datasets. Finally, compared to other CRW datasets, intronic rRNA markers are closest to the simulation study model conditions, but still not exactly the same. The former involve secondary structure evolution, strong selective pressures, and other evolutionary and biophysical constraints that are not accounted for by the estimation/re-estimation methods in our study. This plays a role in the somewhat lower PR-AUC values observed in the empirical study, as compared to the simulation study.

## 5 Discussion

Our findings reflect some basic observations about random walk resampling. The neighbor preservation property is a critical difference between RAWR resampling and bootstrap resampling: the former has it, while the latter does not. The neighbor preservation property helps to ensure that meaningful sequence homology is retained within a resampled replicate and subsequent alignment/tree re-estimation is well-defined.

We note that, compared to SERES, RAWR is closer in design to the bootstrap. There is no need for an anchor selection step and its additional parameters. A simpler formulation should be more tractable to both experimental and theoretical study.

Random walk reversals are essential for producing more replicates beyond the two possible via mirrored inputs, and increased resampling generally improves support estimation. However, the increased resampling replication comes at a price. Reversal introduces a form of “noise injection”. Ideally, if perfect sequence alignments were attainable, all nucleotides near a breakpoint would be correctly aligned and reestimated homologies would be “synchronized” in terms of sequence homology; in practice, incorrect subsequence alignments near a reversal breakpoint introduce the possibility for “de-synchronized” re-estimations (i.e., aligning nucleotide pairs for which homology is not well defined). Too low of a reversal probability effectively limits the number of distinct resampled replicates that are possible, but too high of a reversal probability results in more “noise” that can impact downstream re-estimation. Type I/II error of downstream support estimates are somewhat impacted by this choice, although experiments suggest that a reasonable choice is to err on the smaller side for *γ* settings.

There is another connection between RAWR resampling and bootstrap resampling. Higher *γ* values also have the effect of reducing sequence dependence in RAWR replicates. Lack of sequence dependence is a primary characteristic of bootstrap resampling. These assertions can be seen via some thought experiments. RAWR with *γ* = 0 is equivalent to mirrored inputs but with a random start point and reflection at start/ends of sequences. As discussed in (Wang *et al.*, 2018), RAWR with *γ* ≈ 0.5 is a first-order Markovian process, and RAWR with *γ* < 0.5 requires a higher *r*th-order Markovian process, with smaller *γ* requiring larger *r*. The order *r* can be used to quantify gain/loss of sequence dependence in resampled replicates. We view *γ* ≈ 0.5 to be very large and, as mentioned above, smaller values are likely to be more practical for most applications. (As an aside, RAWR with *γ* = 1 is equivalent to sampling a single site many times and should result in catastrophic data loss.) Results from our experiments with increasing RAWR reversal probability *γ* are consistent with this thinking.

Finally, if the annotation MSA was perfectly accurate, the stationary distribution of a suitable *r*th-order Markovian process would become equivalent to the bootstrap (in the limit of sampled sequence length). But annotation MSAs are never totally correct in practice. Therefore, RAWR’s performance improvement must be due in part to annotation MSA inaccuracy and data augmentation using RAWR replicates, where re-estimation effectively explores more of the underlying search space for the MSA and tree estimation problems.

## 6 Conclusion

In this study, we introduced RAWR, a new non-parametric resampling technique. We applied RAWR to the widely studied task of phylogenetic support estimation. On simulated and empirical benchmarks spanning a range of dataset sizes and sequence divergence, we found that RAWR support estimates had comparable or often better type I and type II error compared to other state-of-the-art methods. Experiments using 10- taxon datasets demonstrated that the improvements were robust to the choice of estimation/re-estimation method and RAWR reversal probability *γ*. RAWR resampling/re-estimation was faster than GUIDANCE2 – a purpose-built parametric method – but slower than a standard bootstrap-based pipeline. The tradeoff between accuracy and computational runtime can be offset through reduced resampling replication.

We conclude with thoughts on directions for future research. First, model misspecification in the empirical study can be ameliorated by including statistical methods to perform inference under more highly parameterized RNA sequence evolution models. Second, orthogonality of non-parametric and parametric resampling techniques brings independent value. Just like in our earlier 2018 study of MSA confidence intervals (Wang *et al.*, 2018), the two classes of resampling approaches can be combined to yield further performance enhancements. Additional improvements may be obtained by combining RAWR resampling with parametric MSA resampling techniques. Finally, non-parametric resampling techniques like RAWR typically require fewer assumptions and are not defined on or constrained to a specific application, unlike parametric and semi-parametric resampling methods. RAWR can therefore be easily applied to any problem with sequence inputs or sequential dependence. We envision many future applications in computational biology and bioinformatics and beyond. Examples include statistical inference of species trees under non-i.i.d. models of sequence evolution (Thorne *et al.*, 1992; Wang and Liu, 2016), machine learning using temporal models (Breiman, 1996), and many, many more – far too many to list here.

